# NRF-1 and HIF-1α modulate activity of human VDAC1 gene promoter during starvation and hypoxia in HeLa cells

**DOI:** 10.1101/2020.06.25.168807

**Authors:** Francesca Guarino, Federica Zinghirino, Lia Mela, Xena Pappalardo, Angela Messina, Vito De Pinto

## Abstract

VDAC (Voltage Dependent Anion Channel) is a family of pore forming protein located in the outer mitochondrial membrane. Its channel property ensures metabolites exchange between mitochondria and the rest of the cell resulting in metabolism and bioenergetics regulation, and in cell death and life switch. VDAC1 is the best characterized and most abundant isoform, and is involved in many pathologies, as cancer or neurodegenerative diseases. However, little information is available about its gene expression regulation in normal and/or pathological conditions. In this work, we explored VDAC1 gene expression regulation in normal conditions and in the contest of metabolic and energetic mitochondrial dysfunction and cell stress. The most active area of the putative promoter region was characterized in terms of transcription factors responsive elements both by bioinformatic studies and promoter activity experiments. In particular, we found a predominant presence of NRF-1 together with other transcription factors binding sites, involved in cell growth, proliferation, development and we studied their prevalence in gene activity. Furthermore, upon depletion of nutrients or controlled hypoxia, as reported in various pathologies, we found that VDAC1 transcripts levels were significantly increased in a time related manner. VDAC1 promoter activity was also validated by gene reporter assays. According to PCR real-time data, it was confirmed that VDAC1 promoter activity is further stimulated when are exposed to stress. A bioinformatic survey suggested NRF-1 e HIF-1α as the most active TFBS. Their validation was obtained by mutagenesis and overexpression experiments. In conclusion, we demonstrated experimentally the involvement of both NRF-1 and HIF-1α in the regulation of VDAC1 promoter activation at basal level and in cell stress conditions.

## Introduction

The Voltage Dependent Anion-selective Channels are a small family of transmembrane pore-forming proteins located in Outer Mitochondrial Membrane (1). Atomic Force Microscopy showed that they abundantely cover the membrane, giving it the appearance of a molecular sieve (2). Their peculiar localization and abundance suggests that they play a very important role in traffic control of small soluble metabolites (3), possibly by the implementation of voltage-dependent gating of the pore (4). It also suggests that they can mediate physical contacts between the surface of the mitochondrion and other external organelles and/or cytoskeletal structures (5-7).

In almost any vertebrate organism the presence of more VDAC isoforms has been established (8): these isoforms have very conserved sequences and their structure are apparently similar, at least for what concerns the transmembrane channel, which is a b-barrel (9). Investigations about specific operation of different isoforms are concentrated on subtle distinctions in the sequences and in the organization of the N-terminal domain, containing helical elements, that results to be the most variable part among the isoforms (10-11).

VDACs, as most of the mitochondrial proteins, are encoded by nuclear genes (1). We expect their expression to be coordinated with that of other mitochondrial proteins in order to achieve a balanced organelle biogenesis. The targeting to OMM has been studied and fully understood only in simple eukaryotes such as *S. cerevisiae* yeast (12), where, on the other hand, there is only one isoform constantly active (13).

No comprehensive and systematic study of the mechanisms of gene expression of VDAC isoforms in higher eukaryotes has ever been undertaken. There is only one report describing the general features of mouse VDACs genes promoter organization and activity (14). In this work the TSS was established by RACE experiments or by sequencing cDNA clones and by prediction with programs available at that time. The sequence intended to be the VDAC1 promoter in mouse showed to lack a TATA box and to be GC-rich. Interestingly, they found that putative mouse VDAC1 promoter lacked any significant activity in antisense orientation and was the least active among three VDAC genes (14).

We propose that promoter characterization could reveal hidden rules or biological purposes that have led to evolution of different isoforms. In fact, the inhomogeneity of VDAC isoforms expression was reported. For example, VDAC1 is expressed in all tissues while VDAC3 is peculiarly abundant in male germ tissues (15). VDAC1 and VDAC3 may be non-expressed without providing a lethal phenotype (at least in mouse, 16-17) while deletion or inactivation of the gene for VDAC2 is lethal (18). This indicates that, in addition to the overall gene regulation activated in mitochondrial biogenesis, there must be regulatory mechanisms peculiar to each gene. The elucidation of these features may lead to an understanding of even the most characteristic roles of each isoform.

In this work we begin studying the regulatory elements of VDAC gene expression starting from the most represented protein, VDAC1. An up-to-date bioinformatic analysis based on the most relevant public databases was performed. A list of TFBS was drawn and a correlation with cellular metabolisms and pathways was attempted.

Since VDAC1 expression is crucial in tumour and in neurodegenerative diseases enough to be considered a marker for these pathologies (19-21), we decided to study the activity of the VDAC1 promoter in cell stress conditions, typical of metabolic reprogramming in tumour and neurodegenerative diseases. We thus assayed depletion of some nutrients like glutamine and pyruvate, the lack of growth factors from the serum, and reduction of oxygen. In general, in response to the alteration of energetic and metabolic stimuli that we induced on HeLa cells, VDAC1 gene and its promoter activity were up-regulated. The response of main bioenergetic TFs like NRF-1 and HIF-1α was studied. We highlighted, in this work, the function of VDAC1 gene expression regulation, as a tool used by the cell to react in extreme conditions, thus supporting the importance of VDAC1.

## Materials and Methods

### Plasmid constructs and mutagenesis

The 600 bp genomic region encompassing the transcriptional starting sites (TSS) of the human VDAC1 gene, located on chromosome 5: 133340586-133341185 (GRCh37/hg19), and three 200 bp sub-sequences derived from it, were transferred from pUC57 vector by digestion with KpnI and HindIII into pGL3 basic vector (Promega). The construct h1prom0-pGL3 contained the whole 600 bp genomic sequence (Chr5:133340586-133341185); h1prom1-pGL3 contained a 200 bp genomic sequence (chr5:133340586-133340786) and, similarly, h1prom2-pGL3 (chr5:133340786-133340986), and h1prom3-pGL3 (chr5:133340986-133341185). pcDNA3.0-NRF-1 and pcDNA3.0-HIF-1α constructs were obtained by PCR cloning strategy (22-24). In brief, coding sequences of TFs amplified from HeLa cDNA were modified at terminal ends by adding sequences complementary to pcDNA 3.0 vector and recombinants generated by PCR. Mutations of predicted NRF-1 and HIF-1α response elements (RE) in hVDAC1 promoters, were introduced using RE specific primers (Table 1 and Table 2 in supplementary material) by QuickChange Multi site-directed mutagenesis kit (Agilent) according to manufacturer’s instructions.

### Cell culture

HeLa cells were cultured in DMEM medium supplemented with 10% FBS at 37°C and 5% CO_2_ atmosphere. After 24h, cells were incubated in stress conditions using a medium devoid of one nutrient (w/o FBS, w/o Gln, w/o Pyr) or in 1% O_2_ (Galaxy® 48 R/48 S CO_2_ Incubator (Eppendorf)).

### Cell viability assay

HeLa cells viability was estimated by 3-(4,5-dimethylthiazolyl-2)-2,5-diphenyltetrazoliumbromide (MTT) method. 1×10^4^ cells/well of 96-well plates were seeded and, four hours before ending treatment, 10 μl of MTT (5 mg/ml) were added to cell culture. At the end of incubation, medium was removed and 100 μl DMSO was added for each well. Optical density was read at 570 nm. Percentage of viable cells was calculated compared to untreated cells and normalized to medium alone. Three independent experiments were performed and data were statistically analyzed by one-way ANOVA. A value of P<0,05 was considered significant.

### Analysis of mitochondrial membrane potential (ΔΨ)

2×10^4^ HeLa cells per well were seeded in a 96-well plate. After 24 hours, cell medium was replaced with complete medium (DMEM + 10% FBS) or without a nutrient (w/o FBS, w/o Gln or w/o Pyr). After 24 and 48 hours, cells were loaded with 10 nM Tetramethyl Rhodamine Methyl Ester (TMRM) (Molecular Probes) and incubated at 37°C for 30 min. TMRM accumulates in mitochondria and its signal intensity is a function of mitochondrial membrane potential. Dye accumulation was assessed by using Incucyte S3 platform (Essen BioSciences) with a 20x Objective. The relative intensity of red fluorescent signal was analysed by Incucyte S3 Software (Essen BioSciences).

### Quantitative Real-time PCR

0.6 ×10^6^ cells were plated in 25 cm^2^ flasks. After 24h and 48h of incubation in nutrient depleted media or in 1% O_2_, total RNA was extracted using “ReliaPrep RNA cell mini-prep system” (Promega) according to manufacturer’s instructions. RNA concentration and purity were measured by a spectrophotometer and 2 μg were used to synthesize cDNA by QuantiTect Reverse Transcription kit (Qiagen). Real-time amplification was performed in a Mastercycler EP Realplex (Eppendorf) in 96-well plates. The reaction mixture contained 1.5 μl cDNA, 0.2 μM gene specific primers pairs (hVDAC1, hVDAC2, hVDAC3, β-actin (Table 1) and 12.5 μl of master mix (QuantiFast SYBR Green PCR kit, Qiagen). Three independent experiments of quantitative real-time were performed in triplicate for each sample. Analysis of relative expression level was performed using the housekeeping β-actin gene as internal calibrator by the ΔΔC_t_ method (25). Data were statistically analyzed by one-way ANOVA. A value of P<0,05 was considered significant.

### Western blotting analysis

For western blotting analysis, 0.6 × 10^6^ HeLa cells were plated in 25 cm^2^ flasks. After 24 hours, cells were exposed to nutrient or O_2_ deprivation as previously described for 24 and 48 hours. Cells were lysed and 80 µg of protein extract of each sample were separated using SDS/PAGE electrophoresis and transferred to a PVDF membrane. The membrane was blotted with primary antibodies anti-VDAC1 1:1000 (ab34726, Abcam) and anti-α-Tubulin 1:6000 (T9026, Sigma-Aldrich) at 4°C overnight and after the washing step with a secondary antibody IRDye® 680RD Donkey anti-Mouse IgG 1:20000 (AB_2814912, Li-Cor Biosciences 925-68072). The membrane was scanned using the Odyssey CLx imaging system (Li-Cor Biosciences). Data were analyzed with Image J software (NIH) and protein expression was measured using integrated intensity readings in regions around protein bands. α-tubulin was used as an internal control for normalizing protein loading.

### Promoter reporter assay

HeLa cells were plated at density of 0.3 × 10^6^ cells/well in a 6 wells plate. After 24h, cells were transfected with 800 ng of pGL3 constructs and 20 ng of pRL-TK renilla reporter vector by Transfast transfection reagent according to manufacturer’s protocol (Promega). 24 h after transfection, cells were exposed to nutrient or O_2_ deprivation as previously described and after 48h cells were lysed.

Luciferase activity of cell lysate transfected with pGL3 promoter constructs was detected with the Dual Luciferase Assay (Promega) according to the manufacturer’s instructions. Activity of firefly luciferase relative to renilla luciferase was expressed in relative luminescence units (RLU). Variation of luminescence units of treated samples relative to control, were indicated as fold increase (FI). Data were statistically analyzed by one-way ANOVA. A value of P<0,05 was taken as significant.

### In silico VDAC1 gene promoter analysis

The human VDAC1 sequence used in the work is identified by the accession number NM_003374. A genomic sequence, (chromosome 5: 133340586-133341185 (GRCh37/hg19)) of 600 bp encompassing the identified TSS, 400 bp upstream and 200 bp downstream, was manually retrieved from GeneBank database of NCBI in human genome assembly chr5:133340586-133341185 (GRCh37/hg19) and was analyzed by Genomatix software (Genomatix GmbH). This VDAC1 promoter sequence was analyzed using Gene Regulation module of Genomatix software suite. In particular, *MatInspector TFs* application were used to identify potential binding site for transcription factors (TFBS) in input sequence. The database performes the analysis using a Matrix Family Library for core promoter elements in vertebrates with a fixed matrix similarity threshold of 0.85 (26). *Overrepresented TFBS and modules* applications were used to associate a statistical analysis to transcription factors binding sites and to regulative units composed by two transcription factors called modules.

The assessment of TFBS predicted by MatInspector, and the functional analysis of transcription-controlling elements, such as the CpG island mapping and active chromatin domains was performed also using UCSC Genome Browser (https://genome.ucsc.edu). In order to evaluate the effectiveness of the promoter prediction, the h1prom0 sequence was examined comparing it with experimentally validated promoters of VDAC1 gene generated by Eukaryotic Promoter Database (EPD, https://epd.epfl.ch) (27). TSS assembly pipeline of EPDnew version 004 for *H. sapiens* is available at the custom track of UCSC Genome Browser.

## Results

### Structure of VDAC1 gene: transcripts variants and promoter

VDAC1 is the most abundant and characterized isoform expressed in mitochondrial outer membrane. The human VDAC1 gene sequence is located in the chromosome 5 (GRCh37/hg19). In Fig 1 it is reported the graphical view of the human genome structure including VDAC1 and the surrounding regions, with indication of regulative elements as reported in UCSC Genome Browser. Three most likely core promoter regions were identified in UCSC, indicated as VDAC1_1, VDAC1_2, VDAC1_3, from the most distal to the most proximal one. The most active one from Eukaryotic Promoter Database (EPD) version 004, is VDAC1_1, which encompasses a sequence downstream the TSS. The VDAC1_1 promoter was thus selected as starting point for experimental analysis performed in this work. The figure shows the results of the BLAT input sequence named h1prom0 of 600 bp (133340586-133341185), utilized in this work, aligned with the annotated promoter sequence in EPD. CpG island, levels of enrichment of the H3K4Me1 and H3K4me3 histone marks, and RNA Pol2 ChIP-seq data associated to this region confirm its trascriptional activity. The HMM data of Chromatin state segmentation in nine different cell lines highlight that the region selected for our study has the features of active promoter (bright red), and it is surrounded by segments defined as weak promoter (light red) and others as enhancer (orange and yellow).

**Figure 1.**
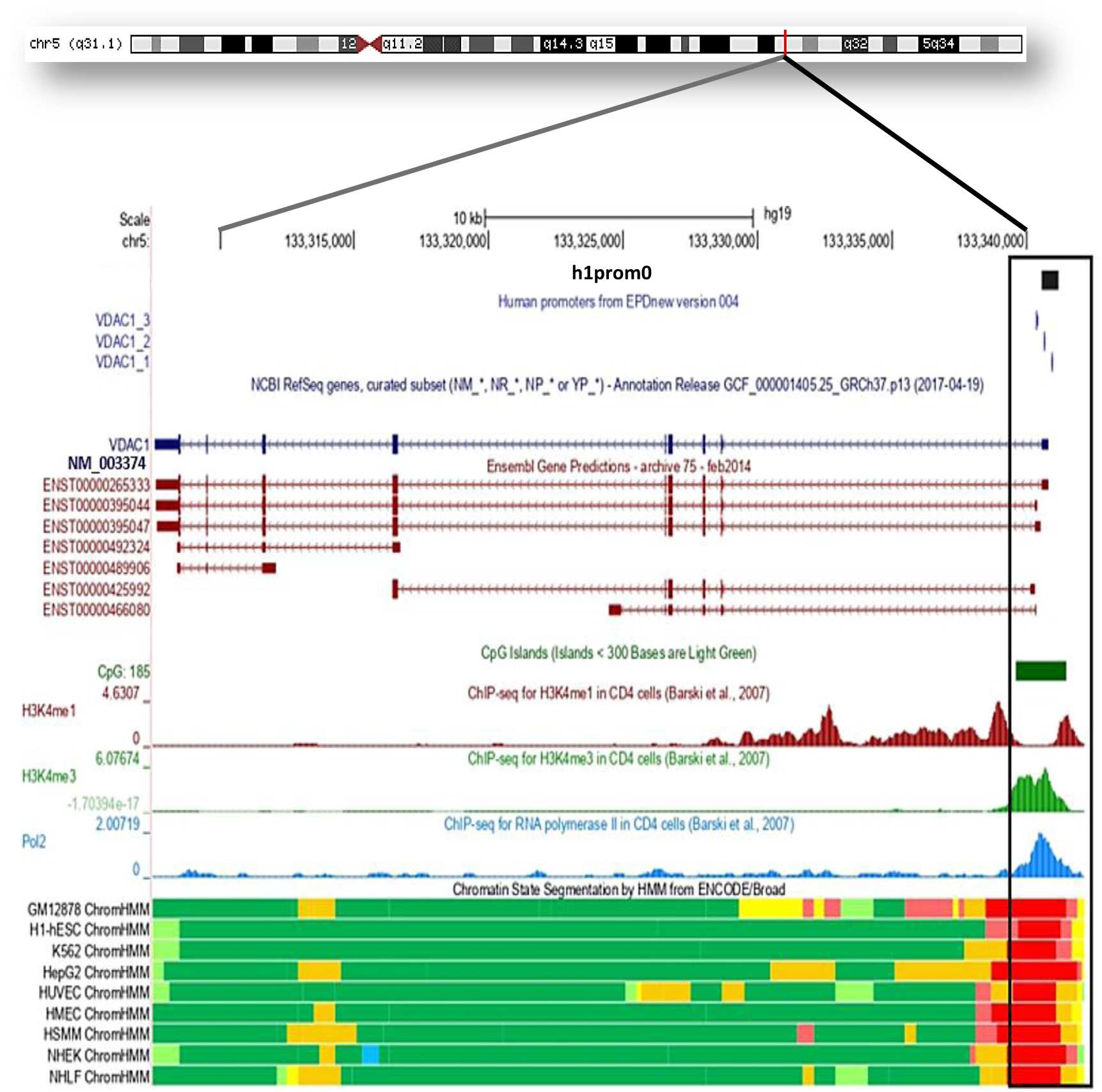
Human VDAC1 promoter structure and function. **(A)** Overview of VDAC1 gene location on Chr5 133340586-133341185 (GRCh37/hg19) obtained in Genome Browser of UCSC. Seven different transcripts of VDAC1 experimentally identified as splice variants were reported with their Ensemble id code. Modified screen shot of *UCSC Genome Browser* shows the results of the BLAT input sequence h1prom0 aligned with annotation tracks of EPD promoter sequences (v.004). Profile of trascriptional activity of this region is highlighted by CpG island identification, levels of enrichment of the H3K4Me1 and H3K4me3 histone marks, and RNA Pol2 and Chromatin State Segmentation ChIP-seq data. The region of trascription activity is overlapping the experimentally identified core promoters reported by Eukariotic Promoter Database (EPD). In the panel are also shown the functional elements highlighted by the data of Chromatin state segmentation by HMM of nine different cell lines which confirm the location of VDAC1 promoter region. Functional elements are identified by different colors as follows: bright red: active promoter; light red: week promoter; orange: strong enhancer; yellow: weak/poised enhancer; blue: insulator; dark green: transcriptional transition/elongation; light green: transcriptional transcribed.

Reported transcripts for VDAC1 are also indicated. They include 7 different splice variants. Four of them encode proteins, two are processed transcripts and in one of them a retained intron is present. Three transcripts have the same exons composition but differ in the lenght of the 5’ and 3’ UTRs. The transcript variant coding the functional protein is reported with the code ENS00000265333.8 corresponding to NM_003374.2 in the refseq database of NCBI. It is not known whether the other splice variants identified have any biological role, however, gene expression data collected from NIH Genotype-Tissue Expression project (GTEX), report the expression of them, including the non protein coding transcripts.

In Fig. 2A the distribution of transcription factors binding sites (TFBS) and general regulatory elements in the h1prom0 promoter sequence was obtained by using Genomatix software (Genomatix GmbH). The statistical significant binding regions were selected by the *Overrepresented TFBS* application. The most represented are V$E2FF TF binding sites, a family of transcription factors involved in cell cycle regulation and proliferation; next V$NRF-1 TF binding sites, that are considered as the master regulator of mitochondrial biogenesis (28, 29). Among the transcription factors binding sites (TFBS) predicted by MatInspector application in h1prom0 promoter sequence, in Fig. 2B are reported TFBS experimentally validated by ChIp-Seq analysis (ENCODE project). The ChIp-Seq Peaks of V$E2FF and V$NRF-1 transcription factors confirm the importance of these factors highlighted by the bioinformatic prediction, suggesting their central role in VDAC1 gene expression regulation. Other binding sites, experimentally validated by ENCODE project data, which are frequently distributed in VDAC1 promoter are V$EBOX, V$EGRF, V$HESF, V$MAZF, V$ZF02, V$RXRF: the corresponding TFs are known to participate in several biological processes, such as cell growth, proliferation and differentiation, development, inflammation and tumorigenesis (30-36).

**Figure 2.**
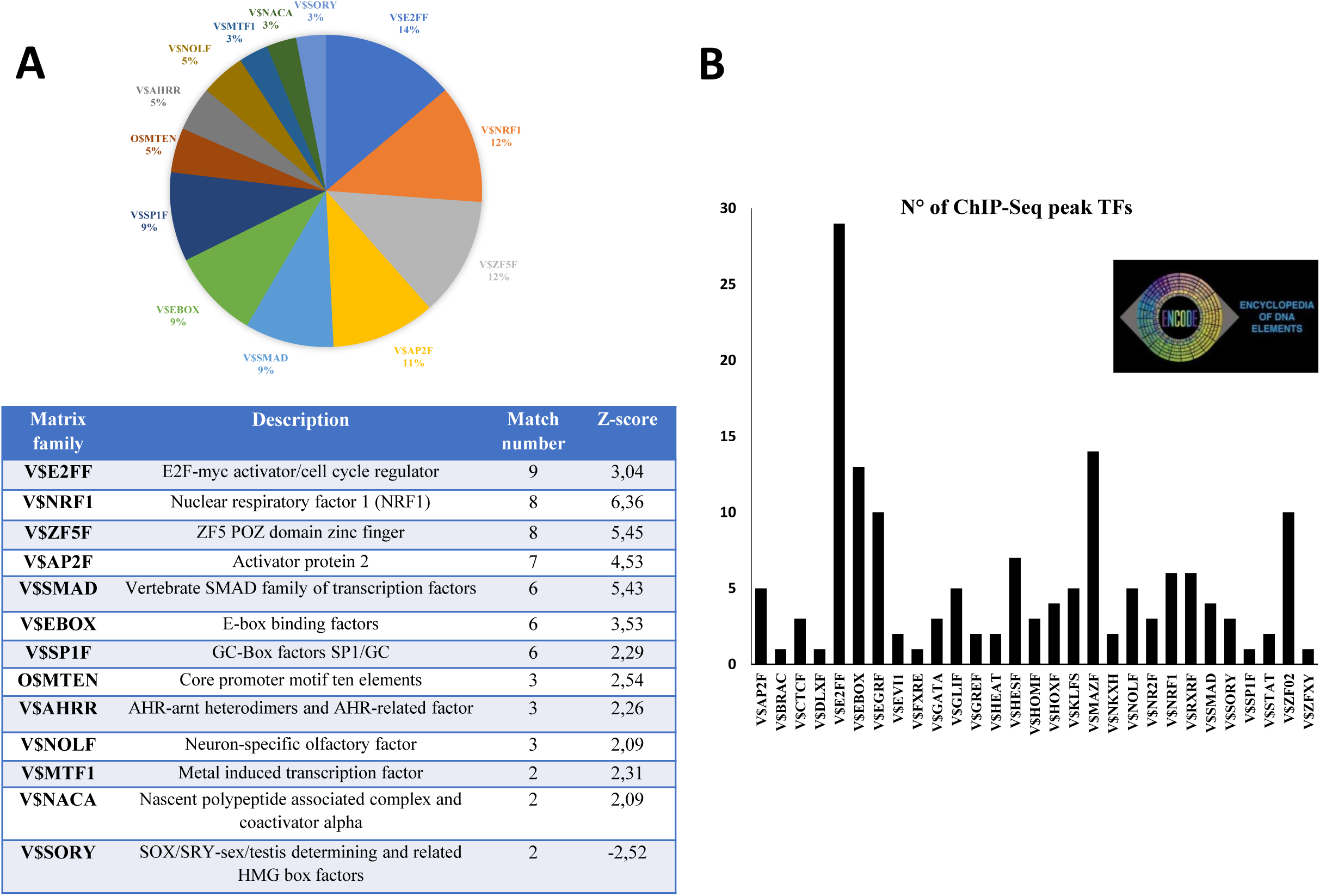
Bioinformatic analysis of human VDAC1 promoter. In panel A it is reported the distribution of Transcription Factors binding sites (TFBS) predicted in h1prom0 promoter sequence by Genomatix software using *MatInspector* and *Overepresented* application. The TFBS reported have been selected by statistical analysis. In the graphical view the percentage of distribution on the sequence analyzed is shown and in the table there are: description of the factors, matches number and Z-score. **(B)** Histogram shows the number of ChIP-Seq peaks for TFBS validated experimentally (ENCODE project) among those predicted by the software Genomatix in h1prom0 promoter sequence.

### Characterization of VDAC1 promoter basal activity

h1prom0 is the genomic sequence of 600 bp encompassing the Transcriptional Starting Site (TSS) cloned and utilized as VDAC1 promoter region for experimental characterization. It was used as a whole or divided in three consecutive parts of 200 bp each. These sequences were cloned in front of the Luciferase (Luc) reporter gene, to study the activity of the human VDAC1 gene promoter. HeLa cells were transfected with pGL3 plasmid containing, upstream the luciferase coding sequence, the whole region of 600 bp of VDAC1 promoter (h1prom0) or three shorter sequences of 200 bp each, h1prom1, h1prom2, h1prom3. Luciferase activity, driven by the whole region of VDAC1 promoter h1prom0, increased 8 folds, compared to pGL3 empty vector (Fig. 3). A similar result was obtained using the 200 bp portion of VDAC1 promoter h1prom1 located downstream the TSS (+1 to +200), suggesting that VDAC1 promoter function is mainly concentrated in this region. On the contrary no activity was reported for the flanking segment h1prom2 (+1 to −200) located upstream the TSS, while the promoter h1prom3 (−200 to −400) next to that, determined a 3 folds activity increase.

**Figure 3.**
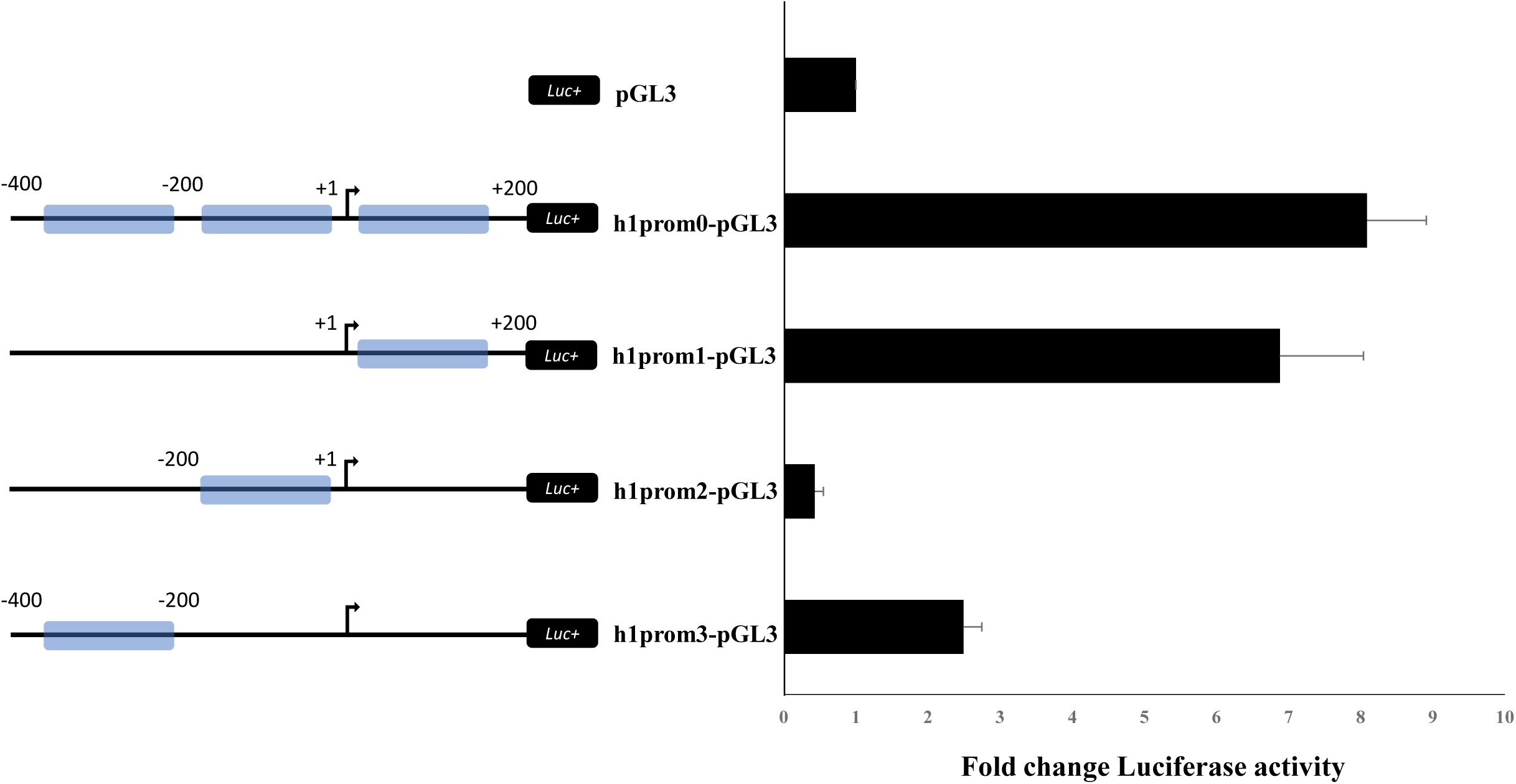
VDAC1 promoter activity. VDAC1 promoter activity was probed by luciferase assay. A 600 bp of genomic region encompassing the transcriptional starting site (TSS) of VDAC1 gene and 200 bp segments of it, were cloned in pGL3 basic vector upward the luciferase gene. HeLa cells were transfected with the constructs h1prom0-pGL3, h1prom1-pGL3, h1prom2-pGL3, h1prom3-pGL3; after 48h cell lysates were prepared and tested for luciferase activity. Fold change of enzyme activity of cells transfected with constructs was reported in comparison to that of cells transfected with empy vector, following normalization with Renilla activity. Data were statistically analyzed by one-way ANOVA. A value of P<0,05 was considered significant.

### Cell viability and mitochondria functionality in normal or stressful conditions

VDAC1 expression and promoter activity was evaluated in HeLa cells in metabolic stress conditions induced by deprivation of nutrients in the medium or oxygen reduction. In this metabolic environment the viability of HeLa cells was tested by MTT assay (Fig. 4A). Cells were incubated with the medium lacking serum, glutamine or pyruvate for 24h, 48h and 72h. Viability of HeLa cells was already reduced by 30% after 24h incubation in any tested conditions. Longer incubations further reduced cell viability: in particular, lack of serum in the medium reduced the cell viability by less than one third after 72h, highlighting the importance of growth factors cocktail present in serum.

**Figure 4.**
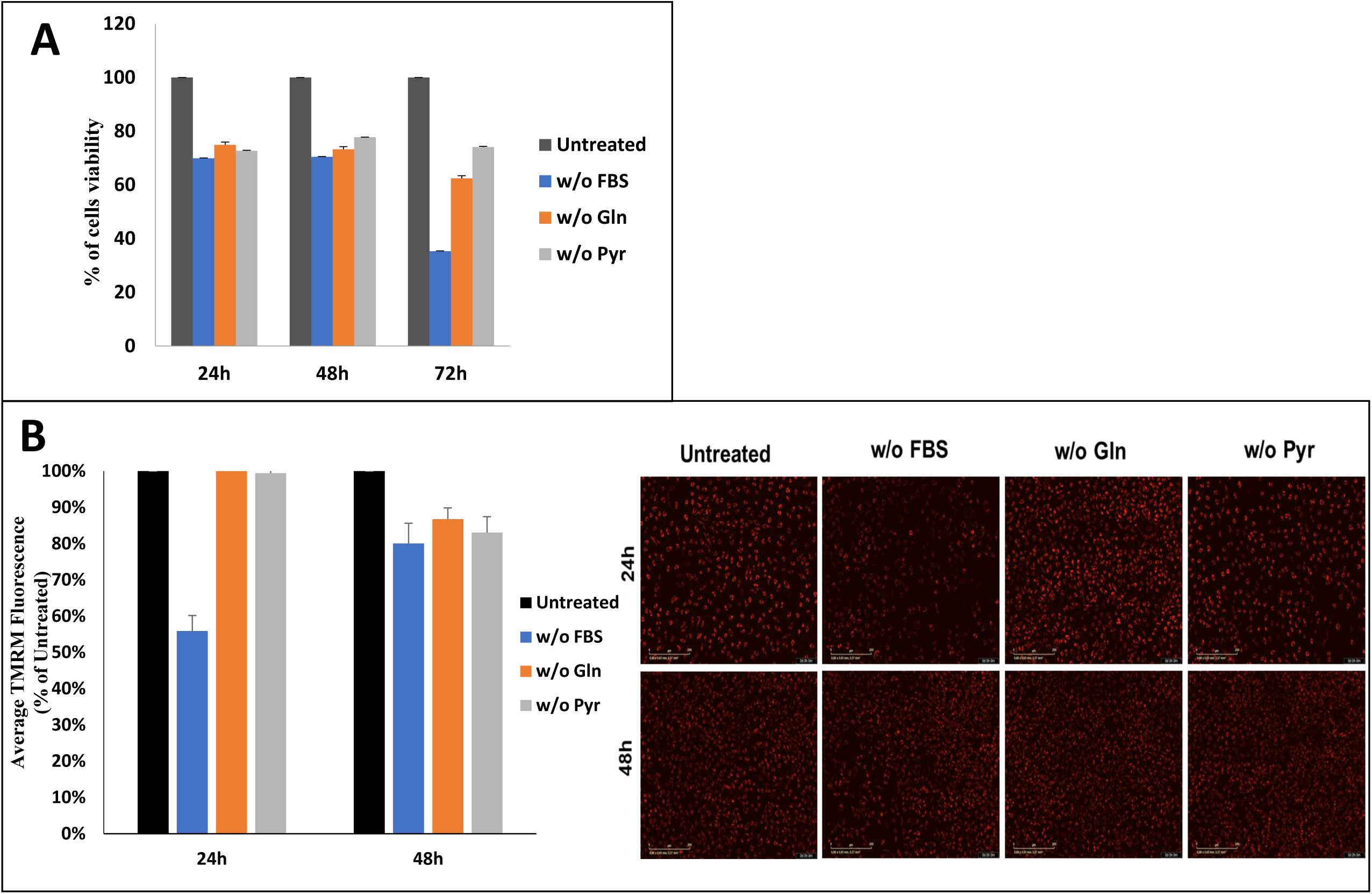
Cell viability and mitochondrial membrane potential in stress conditions. Cells viability and mitochondrial membrane potential was tested in cell depleted of serum, glutamine or pyruvate in the medium. **(A)** Viability of HeLa cells was analyzed at different times (after 24h, 48h, 72h) by MTT assay. Percentage of viable cells, cultured under stress condition, was calculated compared to control and normalized to growth in complete medium. **(B)** Mitochondrial membrane potential of HeLa cells was measured after TMRM probe accumulation in mitochondria in cells under stress conditions (deprivation of serum, glutamine or pyruvate) in comparison to control cells. Dye accumulation was assessed by using Incucyte S3 platform with a 20x Objective and the relative intensity of red fluorescent signal was analysed by Incucyte S3 Software. Three independent experiments were performed for each assay and average results are shown. Data were statistically analyzed by one-way ANOVA. A value of P<0,05 was taken as significant. In the panels on the right side microscopic fields are shown (scale bar 200 µm), with insets showing a magnification of the field. Cells containing TMRM are red.

To assess the effects of nutrient starvation on mitochondrial membrane potential (ΔΨ), HeLa cells were loaded with the mitochondrial ΔΨ-probe TMRM and imaged by fluorescent microscopy. In HeLa cells grown in complete medium for 24 and 48 hours, mitochondria were strongly labeled by TMRM (Fig 4B). After 24h of serum starvation, we observed a 46% decrease of mitochondrial TMRM, followed by the partial rescue of ΔΨ after 48h of treatment. It has been previously observed (37) that the reduction of mitochondrial membrane potential is correlated to decrease of VDAC isoforms expression induced by gene silencing. This result correlates with the fall of cell viability detected by MTT assay. On the contrary, L-glutamine and pyruvate depletion resulted in a slight decrease of TMRM signal compared to the normal growth condition, indicating a minor effect of these deficiencies on mitochondrial membrane potential. Indeed, this result suggests that serum depletion have the major effect on mitochondria and that VDAC1 gene expression and regulation can restore the normal function.

### VDAC1 expression is enhanced in HeLa cells after metabolic stress

There is no systematic and organised information about VDAC1 gene expression and regulation during mitochondrial adaptation to organelle dysfunction, a condition present in many pathologies. VDAC1 mRNA transcription was analyzed in HeLa cells subjected to metabolic stress conditions like serum, glutamine, pyruvate depletion or hypoxia by oxygen reduction to 1%. After 24h a transient effect of VDAC1 gene transcription reduction was obvious but, after 48 h, the transcript synthesis rebounded, increasing at least 2 folds, in comparison to control cells, in absence of pyruvate, glutamine or serum in the medium. The effect due to hypoxia was even more dramatic, since in these conditions the transcript increased almost 5 folds when cells were incubated in 1% O_2_. This result indicates that mitochondria biogenesis is strongly activated (29) (Fig. 5A). The protein levels of VDAC1 were slightly decreased after 24 of nutrient depletion but after 48h of starvation the level is restablished and begins to increase. (Fig. 5B).

**Figure 5.**
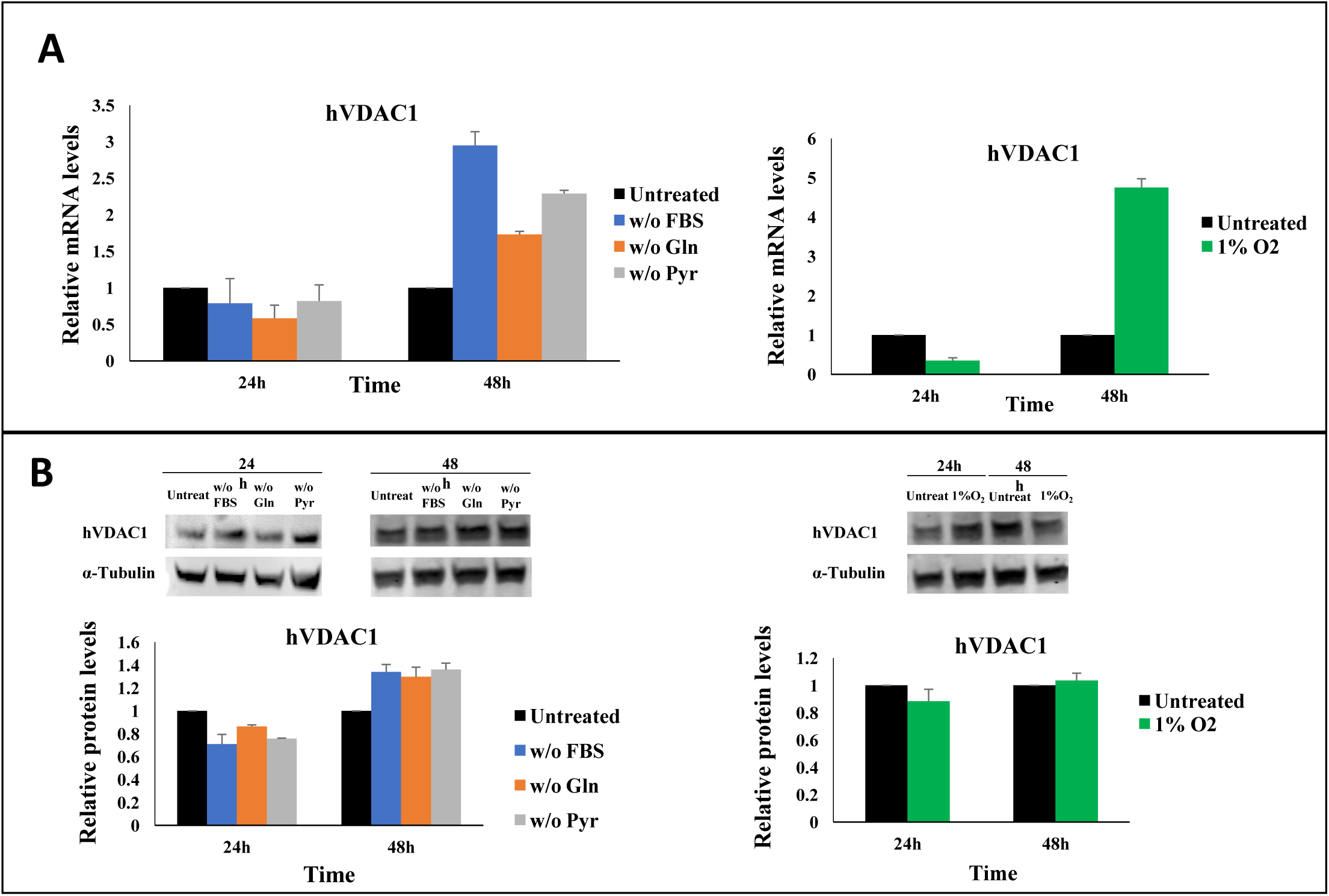
hVDAC1 gene expression and promoter regulation in cell stress conditions. Human VDAC1 expression was studied following exposure of cells to depletion of serum, glutamine, pyruvate in the medium or to hypoxia. **(A)** Transcripts of hVDAC1 were measured, after 24h and 48h of cell treatment, by real-time PCR relative quantification. The variation of hVDAC1 mRNA in treated samples was expressed relative to control samples, after normalization with the housekeeping gene β-actin, by the ΔΔC_t_ method. **(B)** Protein VDAC1 expression was detected by Western blotting of protein from cells incubated in various conditions, after 24h and 48h. Protein extracts were separated by electrophoresis, transferred in a membrane and immunoblotted with anti-VDAC1 antibody (1:1000) and anti-tubulin antibody (1:6000). Anti-mouse secondary antibody IRDye 800CW (1:40000) was used to detect fluorescence by Odyssey equipment.

### VDAC1 mRNA increase in response to stress is due to promoter activation

To explore the VDAC1 mRNA increases following nutrient or O_2_ depletion, the transcriptional activity of VDAC1 promoter (h1prom0, full length) was tested in the same conditions, by measuring promoter activity by Luciferase assays.

In transfected HeLa cells grown in medium lacking serum, up-regulation of 3.4 folds of transcriptional activity of VDAC1 promoter (h1prom0, full length) was discovered. Also the lack of glutamine determined, in transfected HeLa cells, a strong up-regulation of VDAC1 promoter with the increase of luciferase activity of about 10.8 folds in comparison to the basal level. The most impressive increase of luciferase activity (about 18.8 folds) was however detected when cells were grown in hypoxic condition, (1% O_2_) (Fig. 6). In conclusion, the increase of mRNA VDAC1 level is strictly related to response of cellular transcriptional activity when cell culture is modified by depletion of nutrient or in hypoxia.

**Figure 6.**
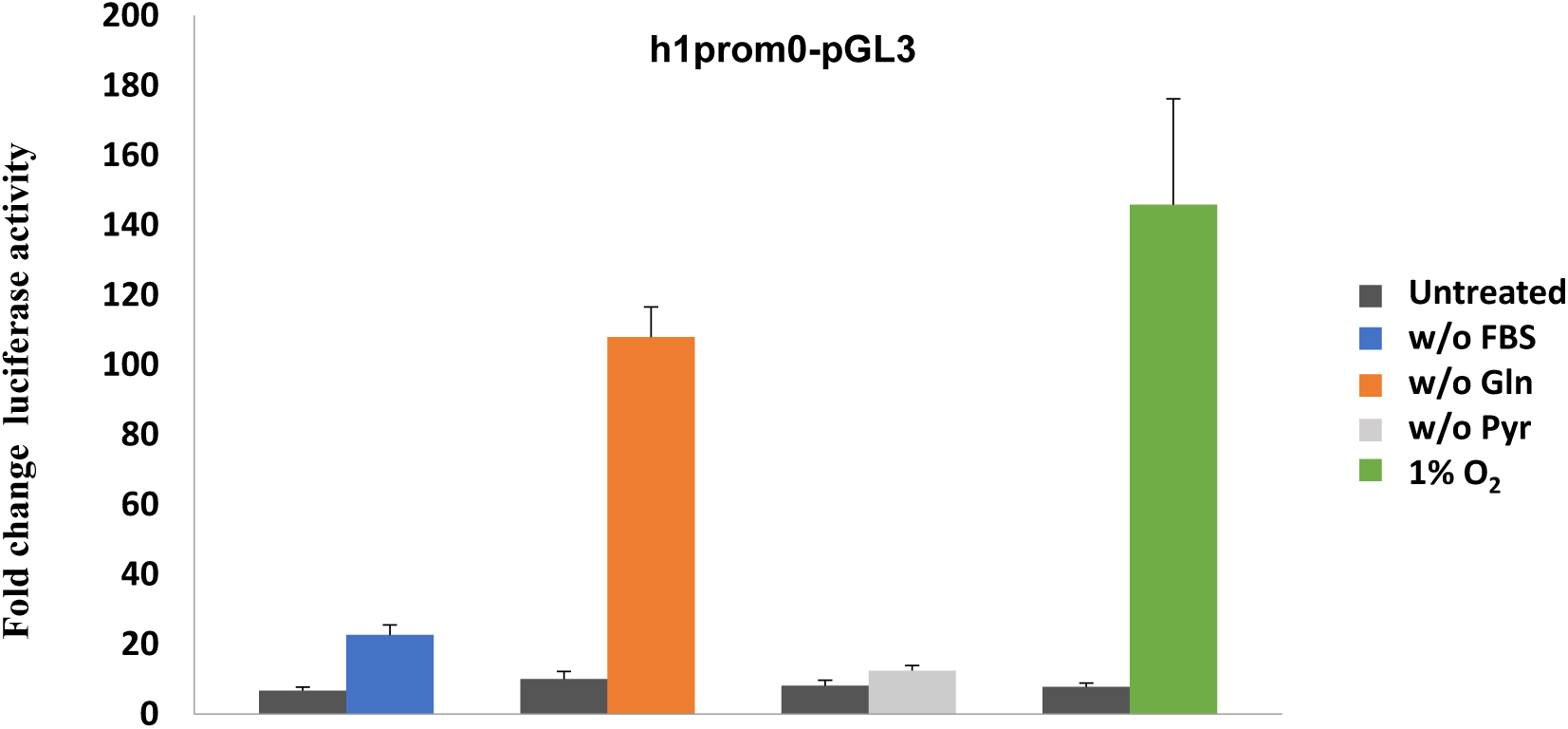
VDAC1 promoter activity in cell stress conditions. Human VDAC1 promoter activity was studied following exposure of cells to depletion of serum, glutamine, pyruvate in the medium or hypoxia. h1prom0 promoter activity was detected by luciferase assay in HeLa cells transfected with h1prom0-pGL3 construct and after 48h of indicated treatments. Luciferase activity of cell lysates was calculated by referring to control cells and following normalization with Renilla activity. Data were statistically analyzed by one-way ANOVA. A value of P<0,05 was taken as significant.

### NRF-1 and HIFF are main regulators of VDAC1 gene expression

We aimed to investigate the molecular mechanism responsible of VDAC1 expression increase in stress condition due to promoter activation we reported above. In particular, we focused our analysis on NRF-1 and HIFF TFs, known to be activated in nutrient depletion and hypoxic conditions (38-40). In Fig. 7A a magnification of VDAC1 promoter region analyzed from UCSC Genome Browser is overlapped with experimental data proving the transcriptional activity of this genomic region, as discussed above in the text. In the same graphical view, the TFBS of NRF-1 in VDAC1 promoter, identified by ChIP-Seq Peaks in ENCODE project, are reported for 6 different cell lines. The two major peaks found for NRF-1 recognition sites are located in overlapping positions of the promoter for any kind of cell. Moreover, these validated TFBS for NRF-1 fall in the genomic region corresponding to VDAC1 promoter sequence studied in this work (h1prom0). At the moment no information in this sequence is available for HIF-1α validated binding sites by the ENCODE project.

**Figure 7.**
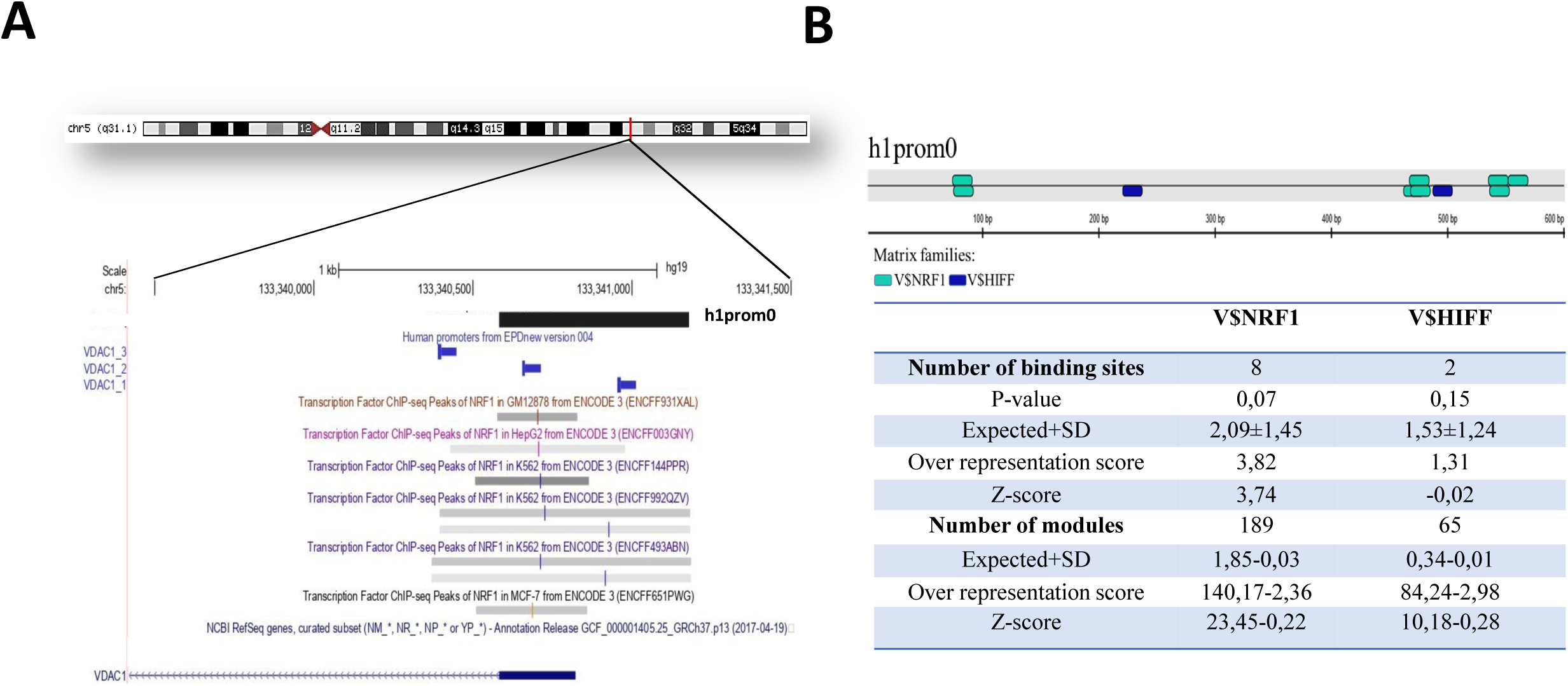
NRF-1 and HIFF transcription factors are regulators of human VDAC1 gene. In (**A)**, a magnified overview of the genomic region including h1prom0 promoter sequence shows the overlapping location of NRF-1 binding sites (bars in the colour segments) identified in 7 different cell lines by ChIP-Seq analysis in ENCODE Project. The horizontal bold line is the 5’UTR sequence joined to the coding sequence (serrated line) of VDAC1 transcript (NM_0033374). In **(B)** bioinformatic analysis of VDAC1 promoter sequence was performed by *MatInspector* and *Overrepresented* applications of Genomatix software. In the graphical view of the sequence, the location of NRF-1 (V$NRF-1) or HIFF (V$HIFF) predicted binding sites in h1prom0 promoter sequence is shown. The number of single transcription factors binding sites (TFBS) of NRF-1 or HIFF and of regulative units (indicated by Genomatix as modules) containing each of the two transcription factors (i.e. V$NRF-1 or V$HIFF) were reported on the table below together with any parameters for the statistical evaluation (p-value, Expected+/- SD, Overrepresentation Score, Z-score).

We also used bioinformatic tools to identify TFBS and regulative modules in VDAC1 gene promoter. *MatInspector* and *Overrepresented* TF applications of Genomatix software allowed to detect the distribution of TFBS. In addition, *Overrepresented modules* application looks for regulative units defined by Genomatix as modules of two clustered TFBS: these modules probably contribute more than the single elements to the promoter activity. 8 NRF-1 and 2 HIFF binding sites were found in the sequence of the promoter h1prom0. We noticed that most of NRF-1 binding sites are located in proximity or over the transcription starting site (TSS), a typical promoter organization found for nuclear genes coding mitochondrial proteins (41-43). It was also reported the number of regulative units, or modules, involving the two considered TFBS: they are 189 for NRF-1 and 65 for HIFF. The whole list of NRF-1 and HIFF TFBS found on VDAC1 promoter is also reported in Fig. 7A, including the results’statistical significance (p-value, expected score, Overrepresentation score and Z-score). These data suggest a relevant role of these factors in VDAC1 gene expression regulation. Moreover, the detection of many modules containing either NRF-1 or HIFF TFBS in VDAC1 promoter suggests that they may have a significant role in the regulation of this gene by promoting a larger cooperation with other TFs in metabolic stress conditions. Analyzing the results of *MatInspector* TF application in details, we found that the two binding sites of HIFF transcription factors family identified on h1prom0 sequence correspond to recognition sites for HIF-1α factor, one of the main regulators of hypoxia cell response.

### NRF-1 and HIF-1α overexpression and mutagenesis of their binding sites confirms their contribution to VDAC1 promoter basal activity

To verify the involvement of NRF-1 and HIF-1α in VDAC1 gene expression, these two transcription factors were overexpressed in HeLa cells where VDAC1 promoter activity was tested. To further validate this result, NRF-1 and HIF-1α TFBS predicted in h1prom0 sequence were mutated (Fig. 8A). Multi-sites concurrent mutations of 5 NRF-1 recognition sites were induced on VDAC1 promoter to generate h1prom0 sequence lacking all of them. With the same approach mutation of 1 HIF-1α binding sites were introduced in h1prom0 sequence. In Fig. 8B VDAC1 promoter transcriptional activity increased up to 11.8 and 9.4 folds when exogenus NRF-1 and HIF-1α factors were expressed, confirming the importance of these two factors in VDAC1 gene expression and regulation, highlighted by the bioinformatic prediction. The transcriptional activity of VDAC1 wild-type and mutated promoters was tested in HeLa cells. The basal transcription activation of VDAC1 promoter was reduced respectively over 2 and 3 folds compared to wild-type promoter when NRF-1 and HIF-1α binding sites were mutated (Fig. 8C).

**Figure 8.**
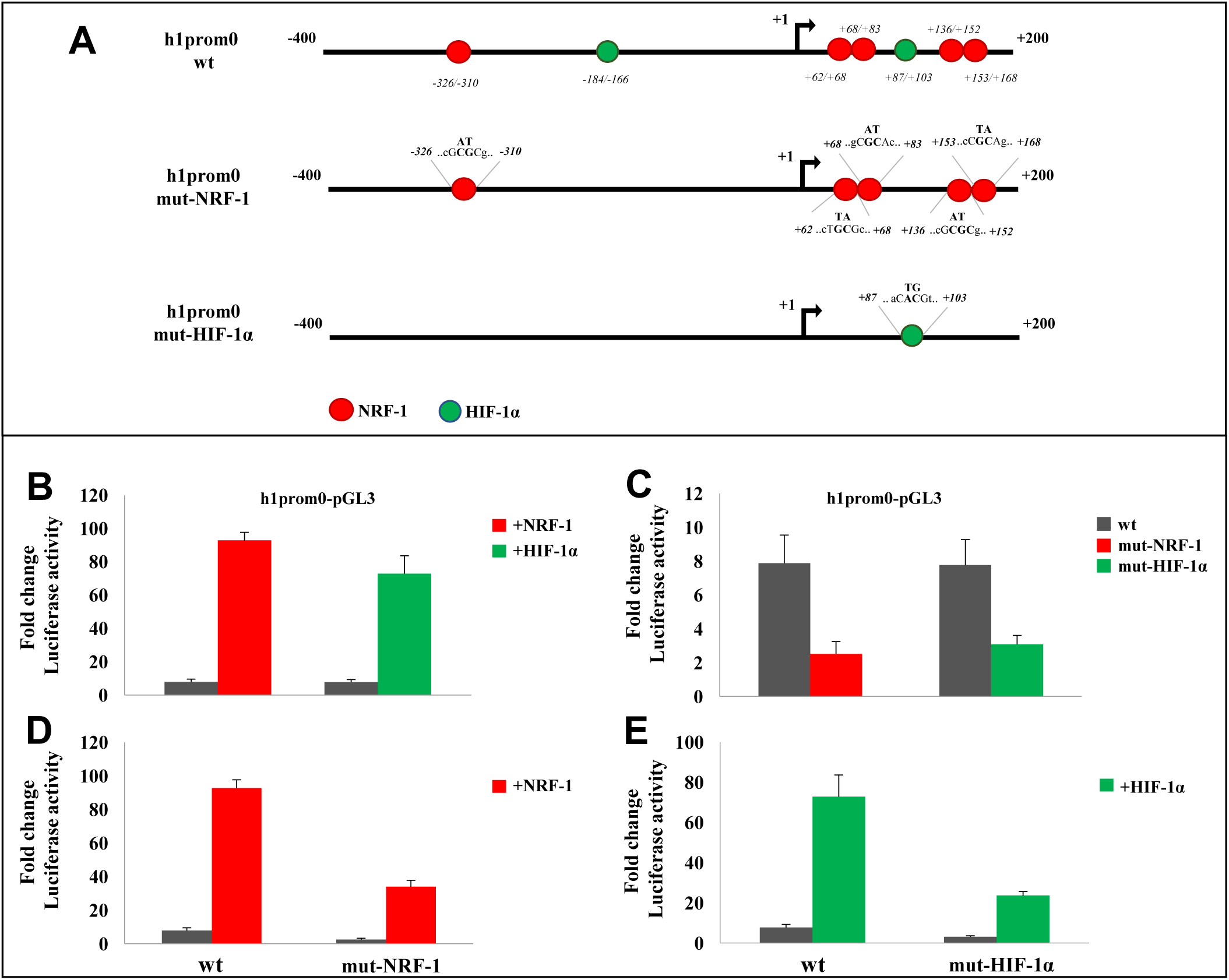
NRF-1 and HIF-1α impact on human VDAC1 transcriptional activity. **(A)** A schematic representation of h1prom0 promoter sequence was drawn with the location of predicted NRF-1 and HIF-1α binding sites. In particular, in h1prom0, the mutated TFs recognition sites were reported highlithing the core sequence and the nucleotides modified for the experimental validation. **(B)** VDAC1 promoter activity was tested by luciferase assay when NRF-1 and HIF-1α transcription factors were overexpressed in HeLa cells. Luciferase activity was measured in lysates of cells, cotransfected with h1prom0-pGL3 + NRF-1-pcDNA3.0 or with h1prom0-pGL3 + HIF-1α-pcDNA3.0 constructs. Trascriptional activity fold change of each samples was reported in comparison to samples transfected with empty vector, following normalization with Renilla activity. **(C)** Luciferase assay performed in VDAC1 h1prom0 containing mutated NRF-1 and HIF-1α binding sites. HeLa cells were transfected with the constructs h1prom0-pGL3, or h1prom0-mut-NRF-1-pGL3, or h1prom0-mut-HIF-1α-pGL3 and after 48h cell lysate were tested for luciferase activity. Trascriptional activity was reported as fold change of h1prom0 promoter in comparison to pGL3 empty vector, following normalization with Renilla activity. In panel **(D)** trascriptional activity of wild-type h1prom0 promoter and the same carrying mutated NRF-1 binding sites, were compared in the presence or absence of ectopic overexpression of NRF-1. HeLa cells were cotransfected with h1prom0-pGL3 wild-type + NRF-1-pcDNA3.0 or with h1prom0-mut-NRF-1-pGL3 + NRF-1-pcDNA3.0. Similarly, in panel **(E)** transcriptional activity of wild-type h1prom0-pGL3 and mutated for HIF-1α recognition sites, was assayed after overexpression of HIF-1α. HeLa cells were cotransfected with h1prom0-pGL3 wild-type + HIF-1α-pcDNA3.0 or with h1prom0-mut-HIF-1α-pGL3 + HIF-1α-pcDNA3.0 constructs. After 48h incubation, cell lysates were prepared and luciferase activity was tested. Trascriptional activity was reported as fold change of h1prom0 promoter in comparison to pGL3 empty vector, following normalization with Renilla activity. Three independent experiments were performed and statistically analyzed by one-way ANOVA. A value of P<0,05 was taken as significant.

In addition, the basal activation of VDAC1 promoter mutated for NRF-1 and HIF-1α binding sites was also evaluated after overexpression of these two factors. As shown in Fig. 8D and in Fig. 8E, the transcriptional activity was reduced up to 2 and 3 folds compared to that of VDAC1 wild-type promoter stimulated respectively with NRF-1 and HIF-1α transcription factors. Therefore, mutations of NRF-1 and HIF-1α recognition sites on VDAC1 promoter drastically compromise the ability to interact with the DNA sequence. The slight rebound of activity of mutated promoters in the presence of transcription factors, in comparison to basal level, can be due either to a reduced but conserved binding to the TFBS, since the mutagenesis affected only two bases in the elements, or to the recruitment of the considered TFs by other factors in the cell, outlining the complexity of the promoters activity regulation (44).

### Role of NRF-1 and HIF-1α on VDAC1 promoter activation in metabolic stress conditions

The role of NRF-1 and HIF-1α was also evaluated in HeLa cells transfected with VDAC1 wild-type or mutated promoters (h1prom0 or h1prom0-mut), and cultured in medium lacking serum, glutamine or in hypoxic conditions at 1% O_2_.

Figure 9A shows the activity of VDAC1 promoter upon overexpression of NRF1 in metabolic stress, i.e. in the absence of serum or glutamine. The activity of VDAC1 promoter was enhanced 3 folds when cells exposed to lack of serum were transfected with vector allowing the overexpression of NRF-1 (Fig. 9A). In the absence of glutamine in the medium, the promoter activity was about five times higher than in the absence of serum. The overexpression of NRF-1 did not stimulate any further increment of VDAC1 promoter transcriptional activity, an indication that the metabolic influence of glutamine requires a different gene activation framework. This result supports the notion that the lack of the cocktails of cytokine and growth factors in the serum has a large influence on the VDAC1 expression.

**Figure 9.**
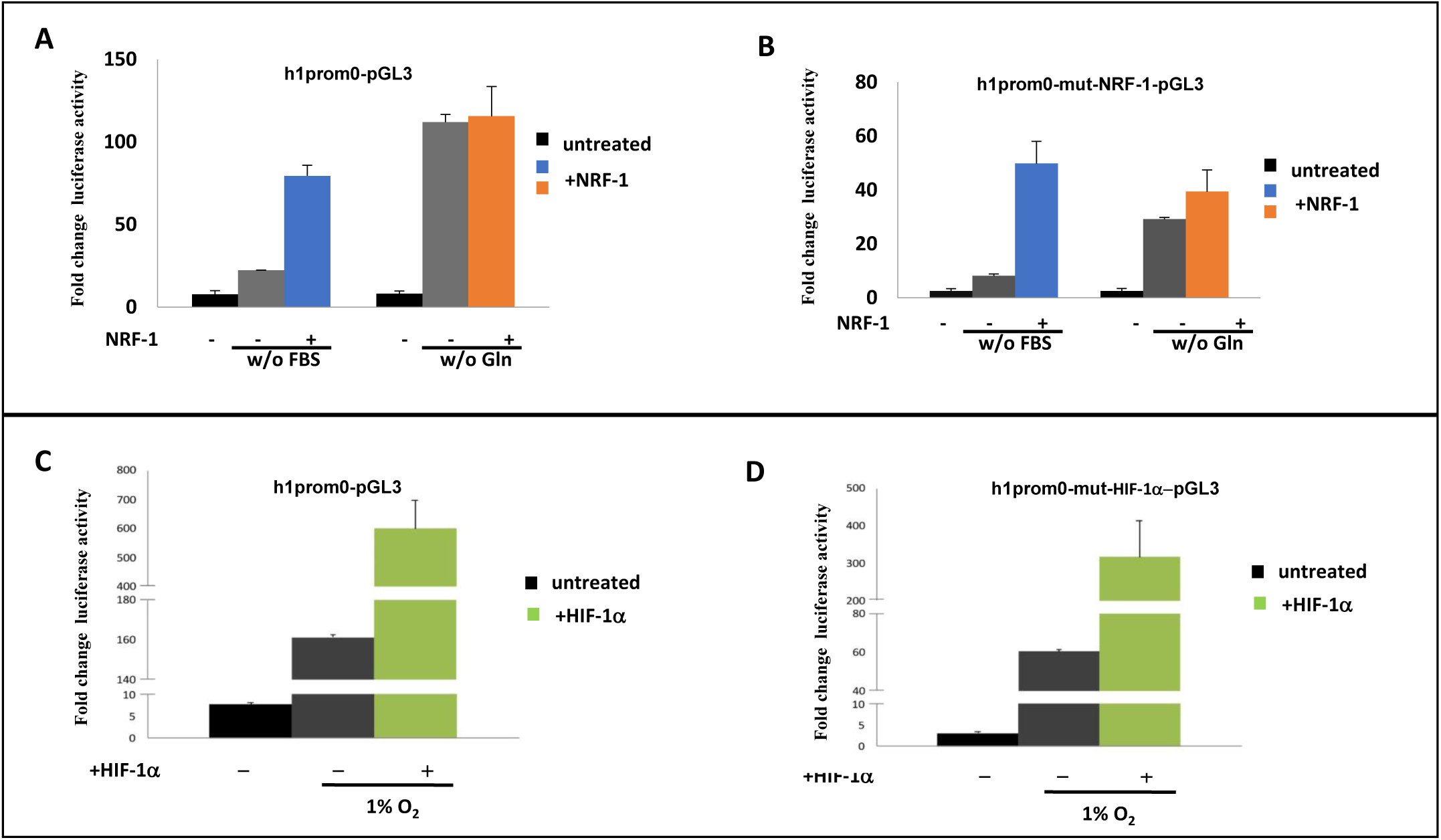
Role of NRF-1 and HIF-1α on hVDAC1 promoter regulation in cell stress conditions. Luciferase activity of h1prom0 wild-type and mutated for NRF-1 and HIF-1α binding sites was assayed in depletion of serum, or glutamine, in the culture medium, or in 1% O_2_ hypoxia. To assess the effect of NRF-1 HeLa cells were cotransfected with the constructs h1prom0-pGL3 wild-type + NRF-1-pcDNA3.0 or with h1prom0-mutNRF-1-pGL3 + NRF-1-pcDNA3.0. To assess the effect of HIF-1α, HeLa cells were cotransfected with the constructs h1prom0-pGL3 wild-type + HIF-1α-pcDNA3.0 or h1prom0-mut-HIF-1α-pGL3 + HIF-1α-pcDNA3.0. After 48h in stress conditions, cell lysates were prepared and assayed for luciferase activity. Trascriptional activity was reported as fold change of h1prom0 promoter in comparison to pGL3 empty vector, following normalization with Renilla activity. Three independent experiments were performed and results statistically analyzed by one-way ANOVA. A value of P<0,05 was taken as significant.

Figure 9B shows the same experiment performed on a VDAC1 promoter where all the NRF-1 binding sites were mutated. The activity of the mutated promoter is overall about 50% weaker than the wild type promoter in any tested condition. The overexpression of NRF-1 activates the transcription, especially in the absence of serum: an apparently surprisingly result, which, however, clearly indicates that NRF-1 has a relevant function in the regulation of gene expression, even though additional TFs are surely needed to fully activate the transcriptional machinery. The result could be explained either by the incomplete inactivation of the NRF-1 binding sites, because only two nucleotides per site were changed, or because other important sites require NRF-1 as an essential co-factor. This notion is reinforced by the result obtained in the absence of glutamine in the medium: overexpression of NRF-1 did not stimulate any further increment of VDAC1 promoter transcriptional activity, as for the wild type promoter, confirming that glutamine is involved in a different regulatory circuit. Indeed, as we defined with the in silico analysis, other than NRF-1, VDAC1 promoter is characterized by a large number of recognition sites for E2FF transcription factors, the family of regulators recruited to control cell cycle, grow, proliferation, mitochondrial biogenesis (45).

Similarly, we evaluated the involvement of HIF-1α on the activity of VDAC1 promoter wild-type and mutated for one HIF-1α binding sites in hypoxic conditions. The transcriptional activity of VDAC1 promoter wild-type was enhanced when cells over-expressing HIF-1α were exposed to reduction of oxygen at 1% (Fig. 9C). The activity of the cell carrying the mutated promoter was about halved in any condition tested (Fig. 9D). However, HIF-1α over-expression stimulated the activity of VDAC1 promoter mutated in cell in hypoxia compared to normal oxygen conditions. Neverthless, over-expression of HIF1α stimulated the activity of both promoters at approximately the same ratio. We suppose that either the HIF-1α site still presents in the mutated h1prom0 can partially support the activity or also for HIF-1α a more complex transcriptional apparatus is active and include HIF-1α on other sites.

### Other TFs involved in mitochondrial biogenesis were predicted to contribute to the regulation of VDAC1 gene expression in stress conditions

To extend our view of the cooperation between NRF-1, HIF-1α and other transcription factors in VDAC1 gene regulation in stress, we deepened our bioinformatic prediction analysis on VDAC1 promoter, looking for other TFBS. In particular, we focused our attention to TFs involved in mitochondrial biogenesis, since VDAC1 gene expression should parallel the increase of mitochondrial mass in the cell, a mechanism used to face cellular stress conditions (29). Based on previous information (38, 46), we searched binding sites for CREB, ETSF, SP1F, YY1F and for modules composed by them together with those for NRF-1.

In Fig. 10A a bioinformatic quest, in UCS Genome Browser, showed a zooming into the VDAC1 promoter region analyzed (chromosome 5: 133340586-133341185 (GRCh37/hg19)) with the distribution of TFBS of regulative factors (CREB, SP1F, ETS, CREM, ATF7, NFE2L2, E2F1) involved in mitochondrial biogenesis and identified by ChIP-Seq Peaks data in ENCODE project. Probably the regulation of VDAC1 gene expression driven by NRF-1 might be triggered in combination with other factors when specific cellular mechanisms are activated in stress conditions. In Fig. 10B, the number and location of these factors is reported in VDAC1 promoter structure, showing that most of them are in close proximity to NRF-1 binding sites. A search for modules containing, as mandatory condition, one of the transcription factor involved in biogenesis in combination with NRF-1, was performed and it was found that high statistical significance is associated to regulative units presenting NRF-1/CREB, NRF-1/SP1F and NRF-1/ETSF. These data corroborate the idea that NRF-1 and HIF1a are not the only necessary TFs to regulate VDAC1 promoter under the stress condition studied, but VDAC1 gene is under a more complex control of the expression.

**Figure 10.**
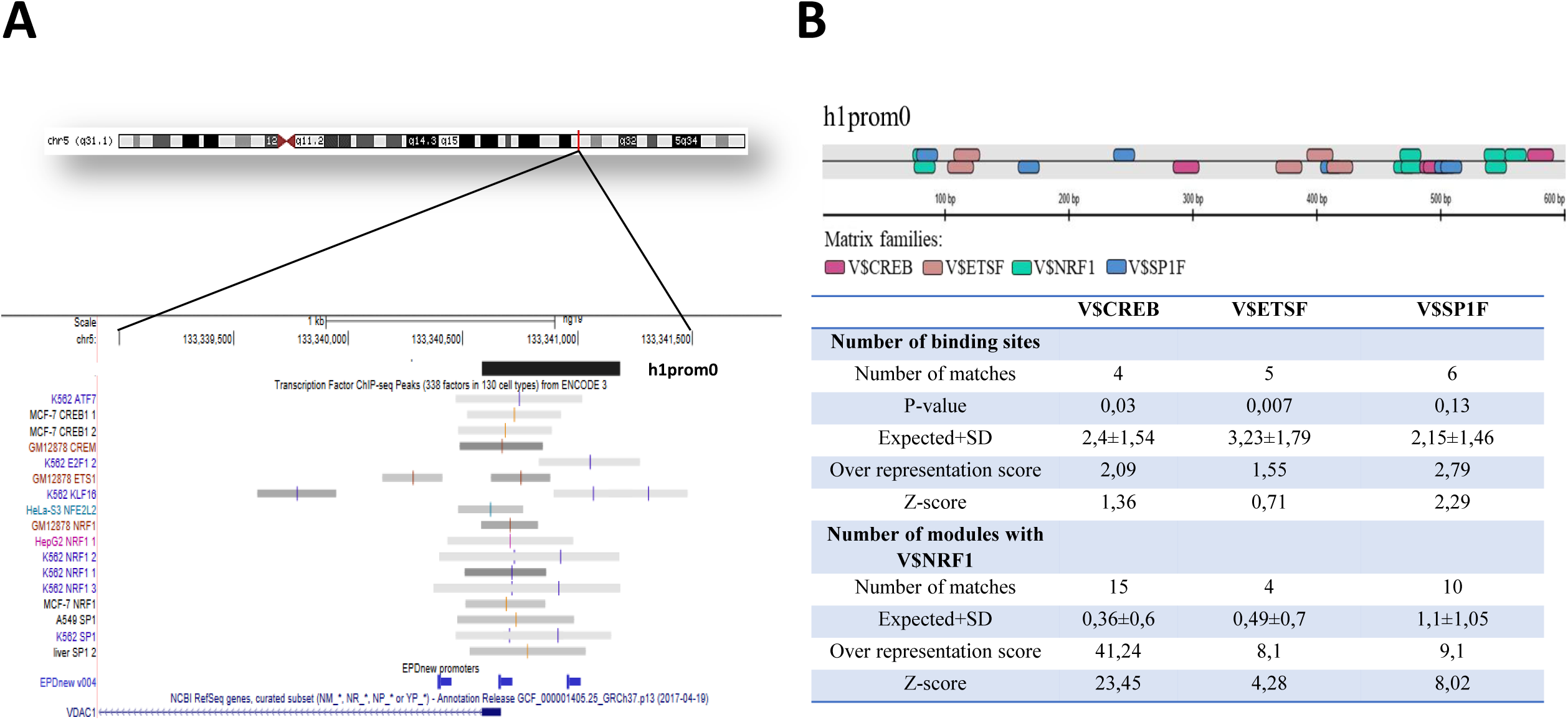
Mitochondrial biogenesis transcription factors in VDAC1 promoter. **(A)** A magnified overview of the genomic region including h1prom0 promoter sequence as in Fig. 3B shows the overlapping location of relevant TF binding sites, involved in mitochondrial biogenesis, (colour vertical bars in the gray segments) identified in different cell lines by ChIP-Seq analysis in ENCODE Project. **(B)** Bioinformatics predictions of single TFBS V$CREB, V$ETSF and V$SP1F and of regulative units composed by two transcription factors (indicated by Genomatix as modules) but including one V$NRF-1, was performed on h1prom0 sequence by *MatInspector* and *Overrepresented* application of Genomatix software. In the graph the location of relevant predicted h1prom0 TFBS, involved in mitochondrial biogenesis, is shown. The number of the single transcription factors and of the regulative modules and any parameter for the statistical evaluation (p-value, expected+/- SD, Overrepresentation Score, Z-score) of V$CREB, V$ETSF, V$SP1F single binding sites and of their regulative modules was reported in the table.

## CONCLUSIONS

VDAC1 is a pivotal protein in the regulation of mitochondrial metabolism and energetic balance in normal conditions but also in tumorigenesis and neurodegeneration. However, the information on VDAC genes expression available in the literature are mainly derived from high-throughput approach and database elaboration data. In particular, it is very common to find correlation of VDAC1 expression changes with proliferative pathways as a main outcome arising from the analysis of many pathologies (47, 48).

We therefore undertook a systematic study of the region upstream of the site where the transcription of VDAC1 mRNA in humans begins, starting with a review of the main public sources and by processing with bioinformatics tools the chosen sequence.

From the in silico analysis of human VDAC1 gene in UCSC genome browser, it appears that 7 different splicing variant mRNAs were identified. This variety does not affect the coding region but mainly modifies 5’-UTR and 3’-UTR length. These sequence maybe target of translation regulation by miRNA or interference. A number of publications indeed reported about the identification of miRNA molecules targeting VDAC1. Among them, miR-7 and miR-320 down-regulate VDAC1 to restore mitochondrial functionality in cellular models of neurodegenerative disease (49) to reduce cell proliferation, migration, invasion (50, 51), or to protect cells from apoptosis in various pathologies (52). To understand and clarify this aspect, an important starting point is to know how and when the expression of these transcripts is regulated. However very little information is available in the literature regarding VDAC1 gene regulation.

We thus cross-checked the public data available on VDAC1 promoter in the Genome browser of UCSC with data extracted by Genomatix suite of software to identify the most relevant transcription factor binding sites (TFBS) located in VDAC1 promoter. In agreement with VDAC1 expression data reported in the literature, we found that the main transcription factors regulators of VDAC1 gene are those involved in proliferation, development, tumorigenesis. As typical of nuclear genes coding for mitochondrial protein, NRF-1 was found to be the transcription factor with the major number of recognition sites and with the highest statistical score in VDAC1 promoter.

### VDAC1 expression in normal and stress conditions

To provide an experimental feedback to bioinformatics, we cloned a 600 bp sequence upstream the mRNA canonical TSS, encompassing the most probable core promoters as predicted in UCSC. From this clone three subclones were derived, each containing about one third of the sequence. The promoter assay showed that the closest and the most distal sequences produced the highest luciferase activity. We decided to compare the promoter activity to stress conditions. We choose nutrient deprivation, in particular medium without pyruvate or glutamine, main substrates of mitochondrial bioenergetics pathways, a medium without serum, thus devoid of growth factors, and an atmosphere containing only 1% O2, i.e. a controlled hypoxia, as the most relevant stress conditions for mammalian cell (53-57).

The analysis of VDAC1 mRNA synthesis demonstrated that VDAC1 transcription was enhanced in any tested stress conditions, after an initial lag. In particular, the answer of VDAC1 gene to hypoxia was dramatic, with a large activation. Also the energization of mitochondria, as detected by TMRM fluorescence, was consistent with the results of Real Time PCR. The protein synthesis showed a delay compared to the transcription, so at 48 h there was an almost identical or slightly increased level than the initial one. This represents a typical adaptation of the cell to mitochondria stress, leading to speeding-up the organelle biogenesis. Detection of promoter activity in the same conditions revealed that it is the promoter activity responsible of the raise of VDAC1 transcription. Hodneland N LI et al. reported that the activation of mitochondrial biogenesis in HeLa cells determines an initial decrease of mRNA levels of NRF-1 target genes, (COX, TFAM)) but, after 24h, the level steadily enhanced during the period of 2-6 days. In parallel the amount of mtDNA, mitochondrial mass, mitochondrial filament/network formation increased corresponding to the activation of AMPK/PGC-1α/NRF-1 axis (29, 58). This pathway typically drives mitochondrial biogenesis. Moreover, it is known that in HeLa cells, short term starvation of glucose, L-glutamine, pyruvate, serum, trigger activation of AMPK nutrient sensing pathways aimed to protecting cells by endogenous ROS production at mitochondrial level (59, 60). The coordination of these pathways is mainly due to PGC-1α and related family members (PGC-1β, PRC), downstream effectors of AMPK (61-63).

### NRF-1 and HIF-1α transcription factors are involved in VDAC1 expression regulation

This evidence prompted us to further investigate on the transcription factors able to trigger VDAC1 gene up-regulation in nutrient deprivation and oxygen reduction. Our attention was focused on the master regulators of mitochondrial biogenesis and hypoxia cell response: NRF-1 and HIFF. NRF-1 is a key transcription factor which mediate nucleo-mitochondrial comunication for elaborating a response to cellular energy demands induced by environmental stimuli as growth factors and nutrient depletion, energy deprivation, oxidative stress (28, 64).

As we described, VDAC1 mRNA expression and promoter activity is up-regulated by the depletion of serum, glutamine or by hypoxia, but at a different extent. While lack of glutamine strongly triggered VDAC1 gene regulation, serum deprivation induced a minor activation of transcription compared to basal level. However, the over-expression of NRF-1 further enhanced VDAC1 promoter activation when cells are exposed to serum depletion, while any/no additional effect was registered on VDAC1 promoter when cells were devoid of glutamine. This result confirms that NRF-1 contributes to transcriptional activation of VDAC1 gene in stress conditions that immediately compromise cells proliferation and mitochondrial functionality, as supported by the data obtained on cells viability and mitochondrial membrane potential. VDAC1 thus might behave as the subset of genes regulated by NRF-1 and targeted by E2F, growth regulatory transcription factor family identified by association of chromatin immunoprecipitation with microarrays data (ChIP-on-chip). Most of them are indeed implicated in mitochondrial biogenesis and metabolism, other than mitosis, cytokinesis (65).

On the contrary, the activation of VDAC1 promoter in glutamine deprivation does not require NRF-1 even though effect on VDAC1 transcription is stronger. In this case we suppose that other transcription factors could be recruited on VDAC1 promoter sequence to activate transcription of the gene in this condition. Therefore, NRF-1 is not the only co-activator of PGC-1, effector of the nutrient sensing pathways, because PGC-1 and other related family members might be involved, in an orchestrated program of gene expression for maintaining the correct energetic, metabolic and oxidation balance: ETSF (NRF-2), PPARγ, ERRα, CREB, SP1, YY1 (28, 46). Information obtained by *in silico* analysis of VDAC1 promoter confirmed that also CREB, SP1, ETSF were predicted to have recognition binding sites in the considered VDAC1 promoter sequence with high statistical score: moreover, they could form regulation modules together with NRF-1.

At last, when we exposed cells to hypoxia a strong activation of the VDAC1 promoter was triggered by HIF-1α over-expression which further enhanced the high activity already registered in oxygen reduction. However, the over-expression of exogenous HIF-1α induced transcriptional activation not only of VDAC1 wt promoter but also of VDAC1 promoter mutated for the HIF-1α binding sites. These data suggest that probably other factors are involved in this pathway and trigger VDAC1 response to promptly adapt mitochondrial functionality to oxygen deprivation. We might suppose that in this condition as well as in nutrient depletion, mitochondrial biogenesis is activated. The trend of VDAC1 transcripts expression, in hypoxic condition, is similar to that observed when cells were exposed to starvation. Most likely, as we suggested above, the expression reduction followed by a quick increase after 24h, occurs when AMPK/PGC-1a/NRF-1 pathway is activated. For this reason, we could hypothesize that also in hypoxia NRF-1 could be involved as a main responsible of VDAC1 gene activation. As already mentioned, this is supported by the finding of a common set of genes regulated by downstream PGC-1α effectors and HIF-1α (66-68).

In this work, for the first time, we highlighted the importance of VDAC1 gene regulation, which is actively modulated in extreme conditions as nutrient depletion or oxygen reduction. In this context, cells activate a response aimed to restoring metabolic pathways and mitochondria adaptation. We demonstrated that VDAC1, the major protein of the mitochondrial outer membrane, might be essential to preserve mitochondria activity during this process.

## Acknowledgments

The authors gratefully acknowledge Dr. Maria Carmela Di Rosa (University of Catania) for establishing western blotting experiments.

## SUPPLEMENTARY MATERIALS

**Table 1.**
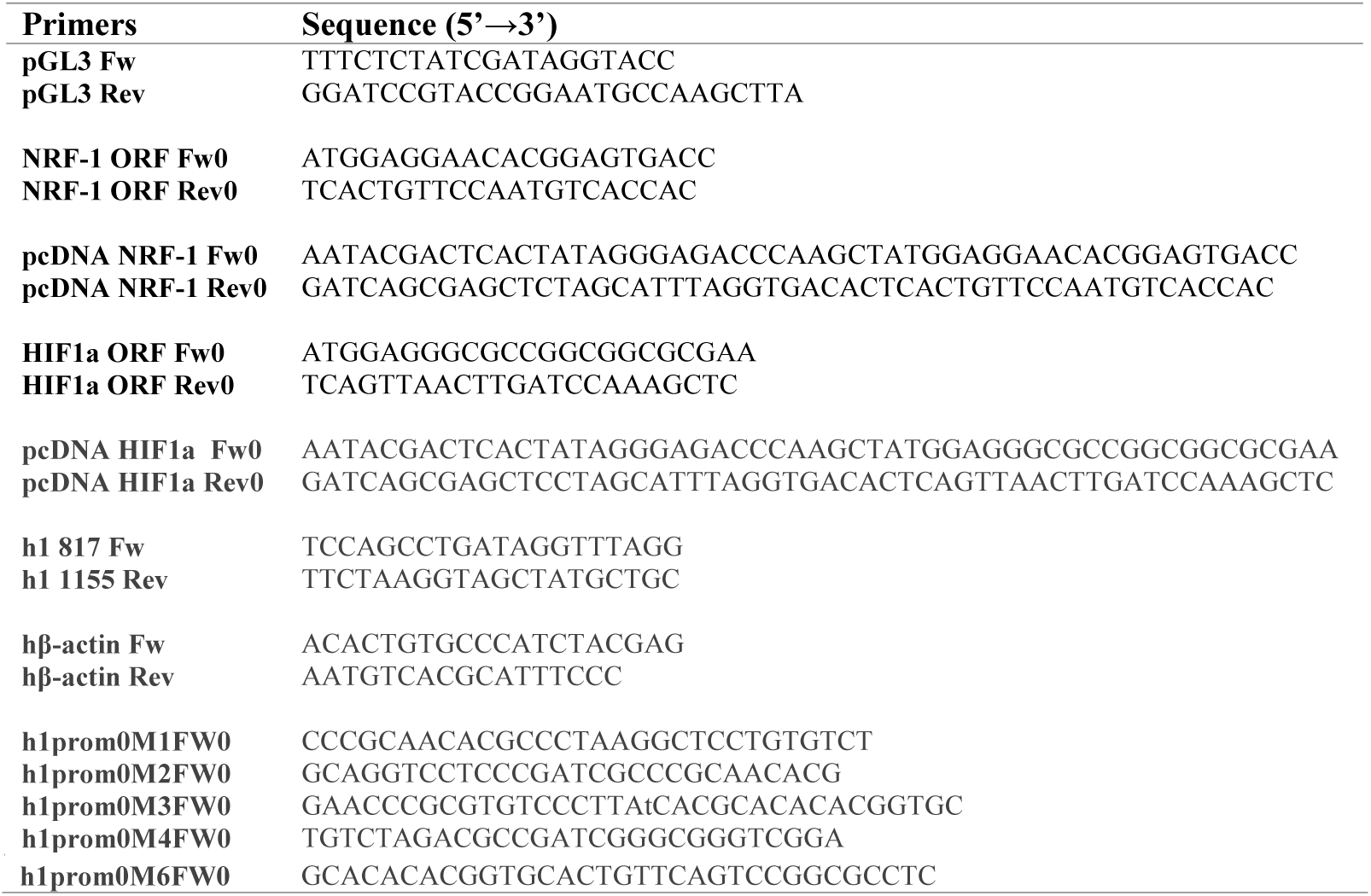

**Table 2.**

| **TF binding sites** | **Genomic position** | **Sequence** | **Mutation** |
| --- | --- | --- | --- |
| **NRF-1** | Chr5:133340659-133340676 | -326/-310:<br>gcc**GCGC**gggcggggtc | M4 : C(-331)>A; G(-332)>T |
|  | Chr5:133341047-133341063 | +62/+78:<br>cgtgtccc**TGCG**cacgc | M3: G(+71)>T; C(+72)>A |
|  | Chr5:133341054-133341069 | +68/+83:<br>ctg**CGCA**cgcacacac | M3: C(+72)>A; G(+73)>T |
|  | Chr5:133341121-133341139 | +136/+152:<br>ccc**GCGC**gcccgcaac | M2: C(+141)>A; G(+142)>T |
|  | Chr5:133341138-133341154 | +153/+168:<br>cgcc**CGCA**ggctcctg | M1: G(+157)>T; C(+158)>A |
| **HIF-1α** | Chr5:133340802-133340820 | -184/-166:<br>gggcc**CACG**cgggcttt | M5: A(-177)>T; C(-176)>G |
|  | Chr5:133341072-133341089 | +87/+103:<br>tgca**CACGT**cagtccgg | M6: A(+92)>T; C(+93)>G |

